## supplementary figures for "The evolution and mutational robustness of chromatin accessibility in *Drosophila*"

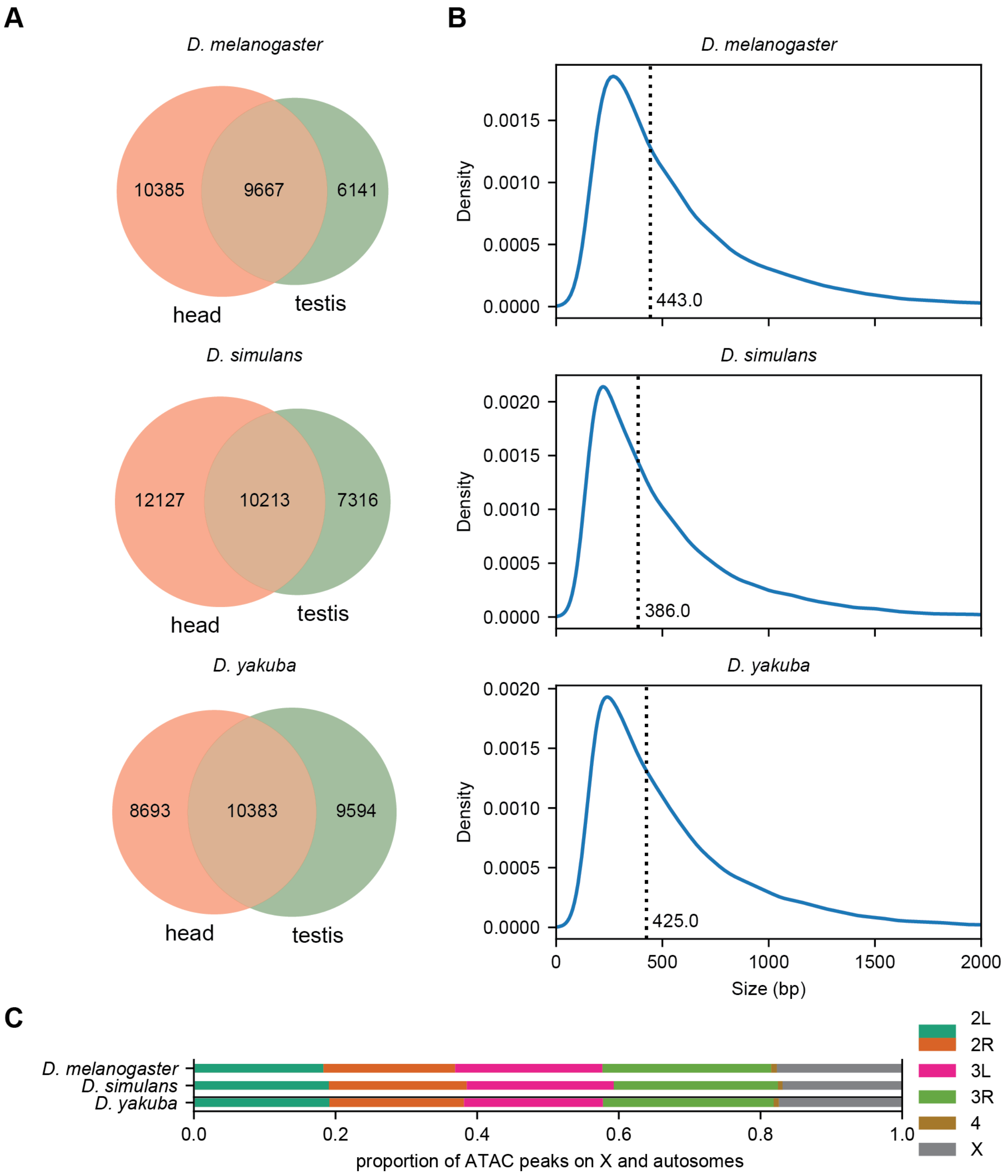


**Supplementary Figure 1. Summary of peak properties.** A) Number of peaks identified in each tissue. B) Distribution of peak size in each species. The dotted line indicates the median. C) Chromosomal locations of peaks. Flies have 3 autosomes (chromosomes 2 and 3, each with L and R arms, and 4) and two sex chromosomes (X and Y). We omit Y because it is very small and gene-poor.

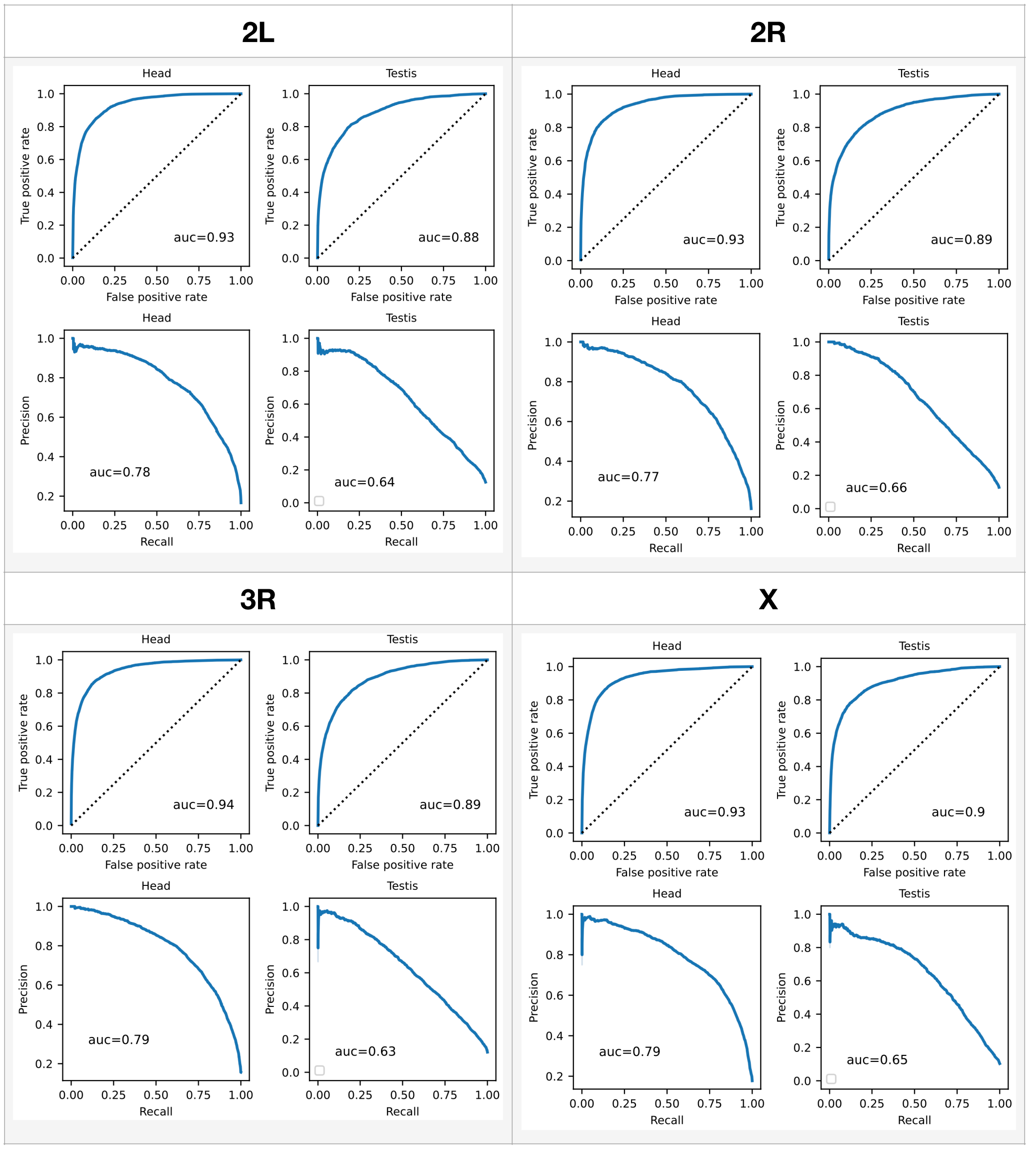


**Supplementary Figure 2. Performance of models on other test chromosomes.** ROC and PR curves for models tested on *D. melanogaster* chromosomes other than 3L. In each case, models were trained (including hyperparameter tuning) on all major chromosome arms except the test chromosome arm.

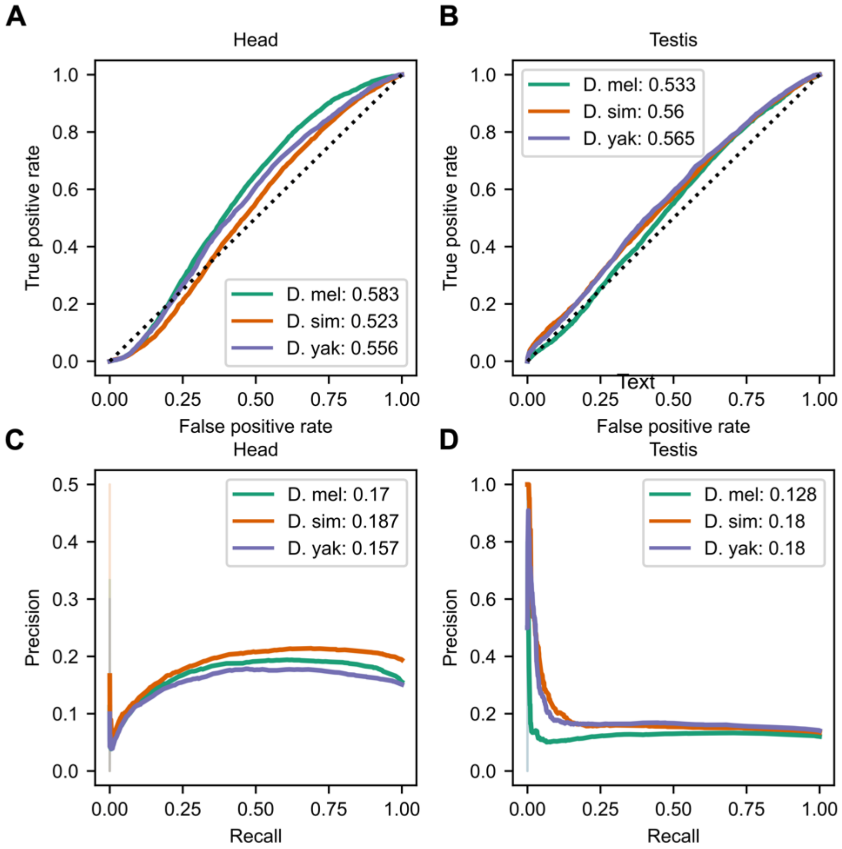


**Supplementary Figure 3. GC content is very weakly predictive of chromatin accessibility**. ROC and PRC curves if the model only learned the GC content of each example. (A) ROC curve showing classification performance in head. (B) ROC curve showing classification performance in testis. (C) and (D) Precision-recall curves showing performance in head and testis respectively.

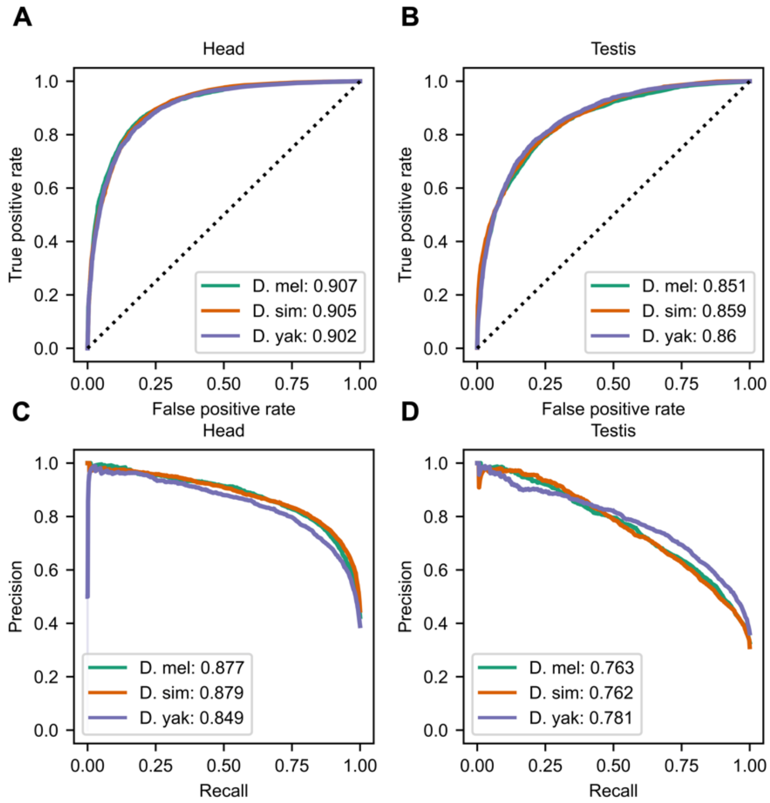


**Supplementary Figure 4. Considering only non-peak examples that are proximal to peak examples does not qualitatively impair model performance**. ROC and PRC curves using non-peak examples that were at most 500 bp away from peak examples. (A) ROC curve showing classification performance in head. (B) ROC curve showing classification performance in testis. (C) and (D) Precision-recall curves showing performance in head and testis respectively.

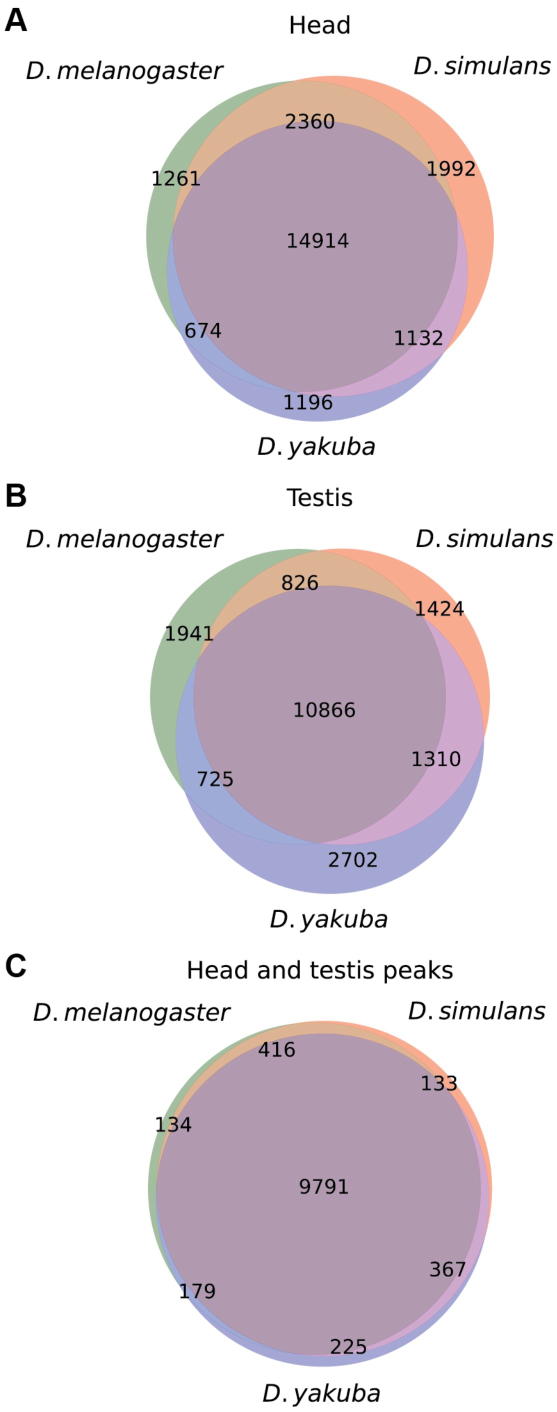


**Supplementary Figure 5. Conservation of peaks across species based on tissue.** A reference-free multi-context approach was used to identify the orthologous regions for each peak in the other two species (see Methods). (A) Venn diagram of peak conservation in the head across species. (B) Peak conservation in testis across species. (C) Conservation of peaks shared in both head and testis across species.

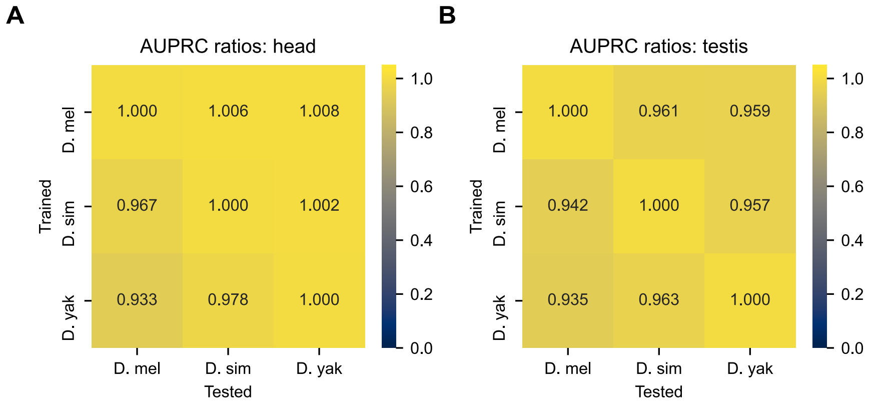


**Supplementary Figure 6. Cross-species performance of models – AUPR ratios.** Each model (trained in a given species) was tested in the other two species. The AUPR of each model tested in a non-native species was divided by the AUROC of the native model.

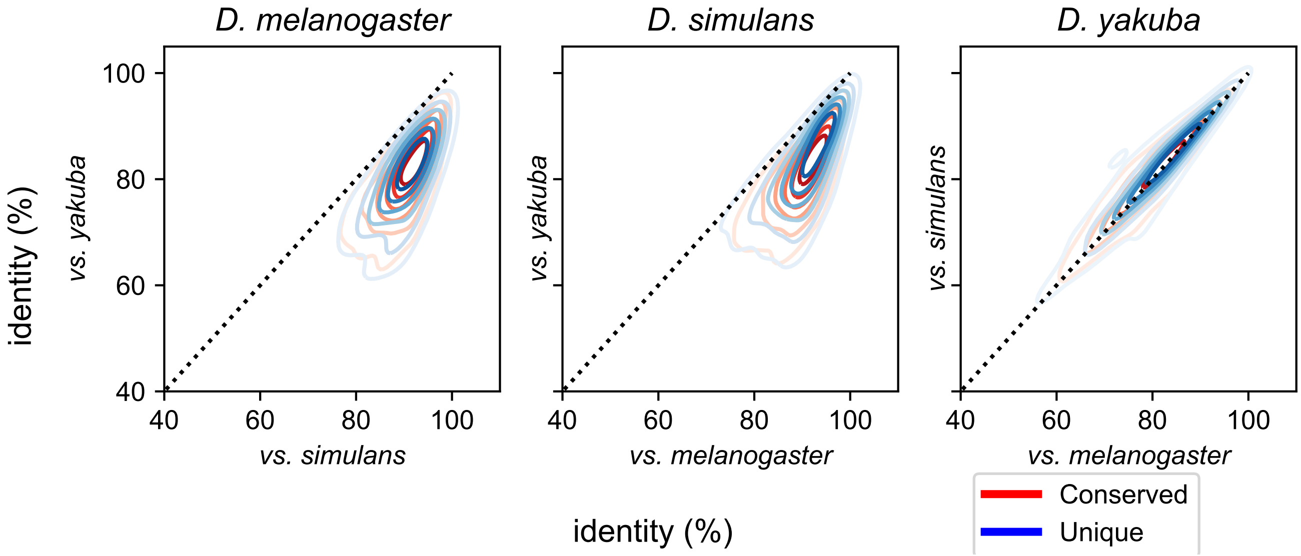


**Supplementary Figure 7. Genomic divergence of peaks.** (A) Kernel-smoothed contour plot of the percent identity of peaks called in *D. melanogaster* versus their orthologous sequences in *D. simulans* (x-axis) and *D. yakuba* (y-axis). Species-specific peaks ("unique"; blue) have similar levels of sequence identity as conserved peaks (red); note generally greater identity with *D. simulans* than *D. yakuba*, consistent with the latter being more distantly related . (B) As in panel A, except with peaks called in *D. simulans* versus their orthologous sequences in *D. melanogaster* ("mel", x-axis) and *D. yakuba* (y-axis). (C) As in panel A, except with peaks called in *D. yakuba* versus their orthologous sequences in *D. melanogaster* (x-axis) or *D. simulans* (y-axis). Note similar identity between comparisons to the two other species here, since *D. yakuba* is the outgroup of the three species.

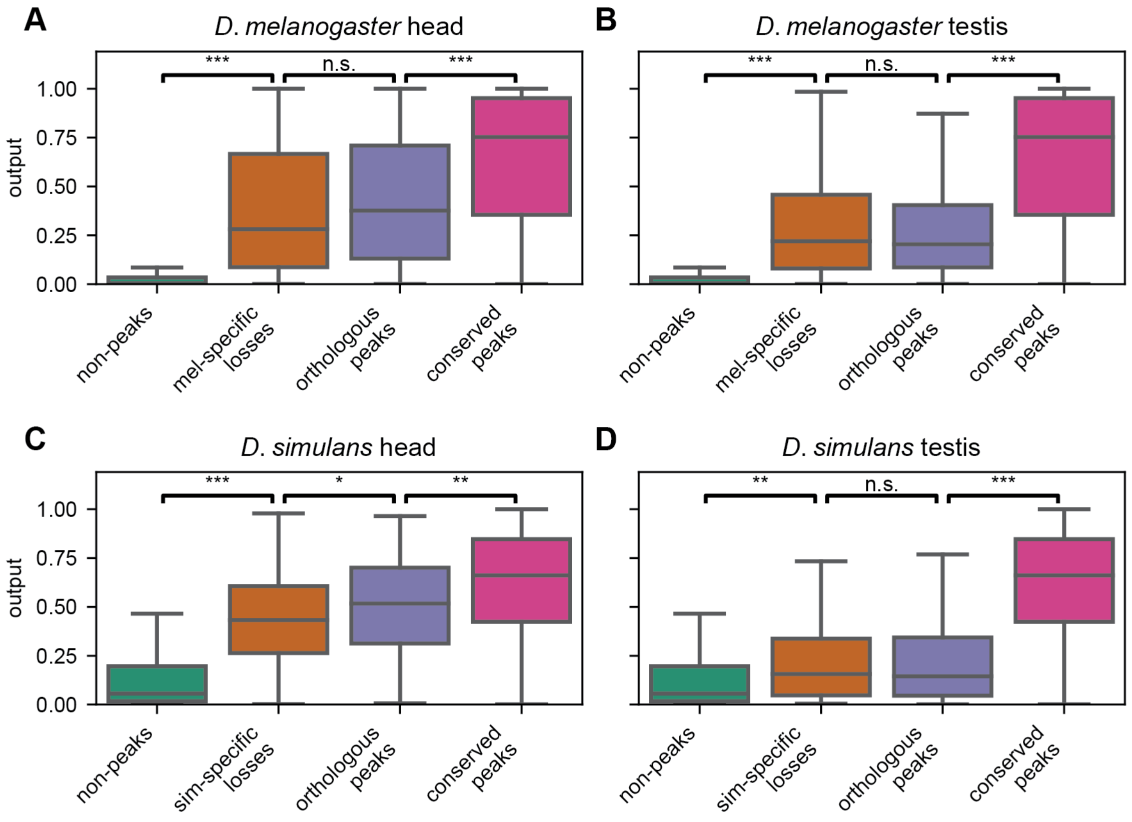


**Supplementary Figure 8. Species-specific losses also show model outputs between peaks and non-peaks.**

Boxplots of raw model outputs, focusing on species-specific losses (that is, peaks with orthologs in *D. yakuba* and exactly one of *D. melanogaster* or *D. simulans*). (A, B) Model outputs for *D. melanogaster*-specific losses were computed for head (A) and testis (B) using the model trained on *D. melanogaster* data from the two orthologous peak sequences (blue; *D. yakuba* and *D. simulans*) and the orthologous non-peak sequences from *D. melanogaster* (orange). Conserved *D. melanogaster* 3L peak examples (magenta) and 3L non-peak examples (green) are shown for reference. The sequences of peaks from *D. simulans* and *D. yakuba* that are orthologous to *melanogaster*-specific losses on 3L (blue) still have dramatically lower model outputs than peaks overall; the corresponding non-peak sequences from *D. melanogaster* (orange) still have dramatically higher model outputs than non-peaks overall. Note that in head, species-specific non-peaks still have lower model outputs, even though they now are from a different species than the species on which the model was trained; in testis, species-specific peaks and non-peaks have very similar distributions. One-sided Mann-Whitney U tests; ***: $p < 1E\text{-}10$; **: $p < 0.001$; *: $p < 0.05$. (C, D) As in panels A and B, except using the *D. simulans* model, the *D. simulans* 3L peak and non-peak examples for reference (magenta and green, respectively), and *simulans*-specific non-peak sequences (orange) and their orthologous peak sequences from *D. melanogaster* and *D. yakuba* (blue).

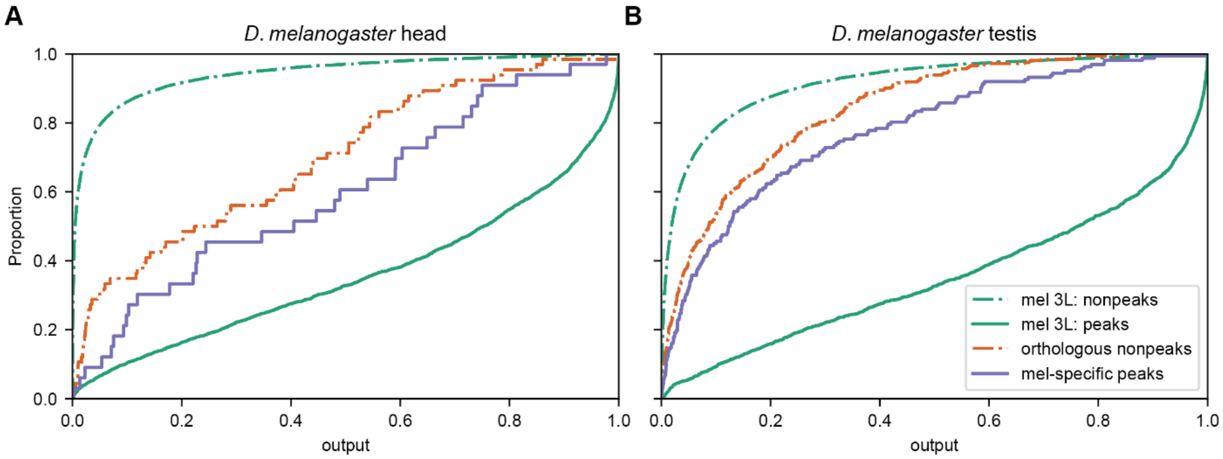


**Supplementary Figure 9. Low coverage sequences orthologous to species-specific peaks show model outputs between peaks and non-peaks.** Cumulative distributions of raw model outputs, focusing only on those species-specific peaks least likely to be falsely-called as species-specific. Cumulative distributions of raw model outputs in head (A) and testis (B) from the model trained on *D. melanogaster* data. Solid green line: 3L peak examples; dashed green line: 3L non-peak examples. High-confidence m*elanogaster*-specific peaks on 3L (i.e., *melanogaster* peaks whose average coverage in the orthologous *simulans* and *yakuba* regions is less than the first percentile of called peaks in both species; solid blue line) still have dramatically lower model outputs than peaks overall; the corresponding orthologous non-peak sequences from *D. simulans* and *D. yakuba* still have dramatically higher model outputs than non-peaks overall. The species-specific peaks have modestly higher model outputs than their orthologous non-peaks.

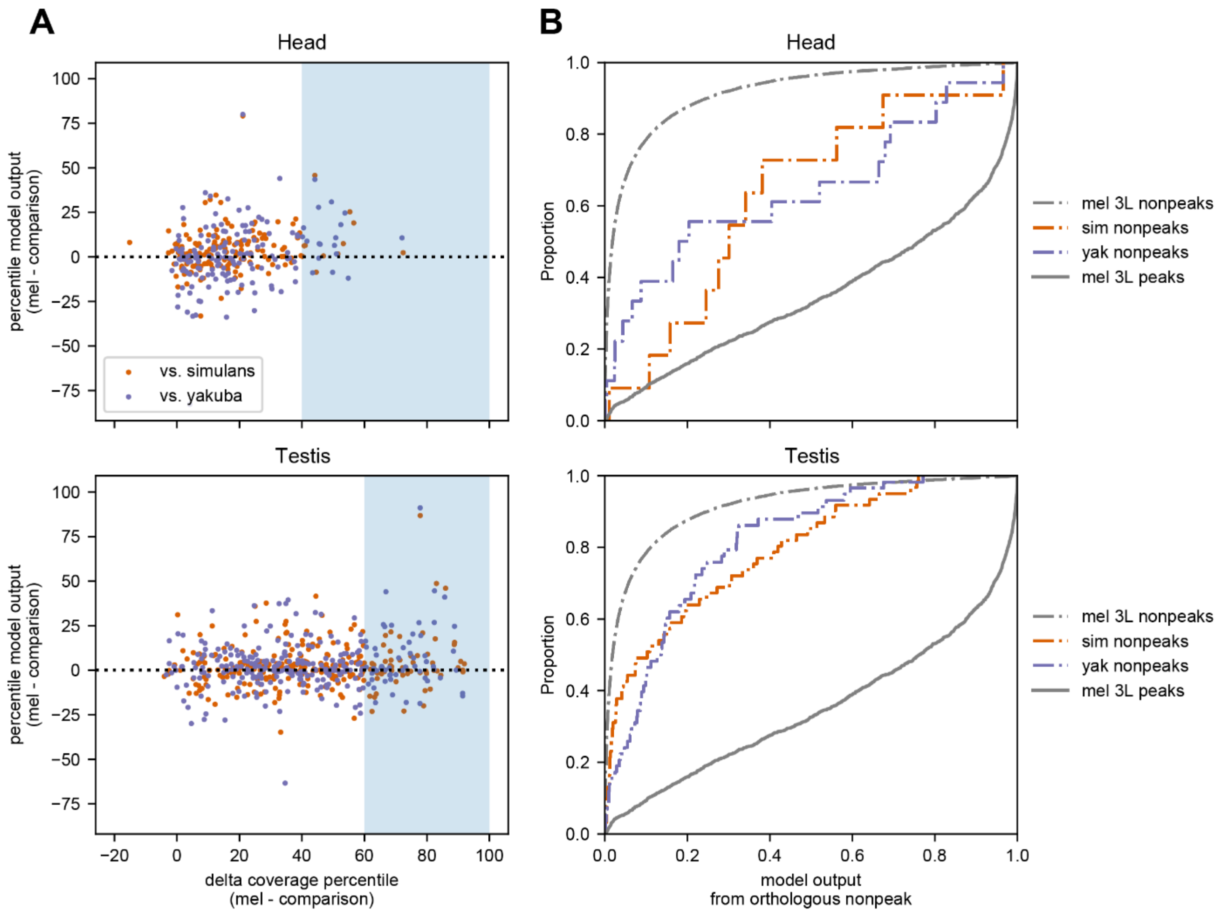


**Supplementary Figure 10. The relative change in model outputs vs. relative change in sequence coverage.**

(A) Considering *melanogaster*-specific peaks on 3L, we calculated the mean ATAC-seq coverage in both *D. melanogaster* as well as the orthologous non-peak regions in *D. simulans* and *D. yakuba*, transformed these into percentile ranks with respect to the respective species (that is, orthologous coverage in *simulans* as a percentile of peaks called in *simulans*), and calculated the change in percentile rank from *melanogaster* to the other species (red: vs. *simulans*; blue: vs. *yakuba*; top: head; bottom: testis). We also calculated the model outputs using the *D. melanogaster* model for both the *melanogaster* peak sequences and the non-peak regions, similarly transformed these into percentile ranks (that is, model output for orthologous sequence in *simulans* as a percentile of peaks called in *simulans*), and similarly calculated the change in percentile rank from *melanogaster* to the other species.

(B) The model outputs of orthologous non-peak sequences with very high relative coverage shifts (panel A, shaded boxes) remain in between those of the *melanogaster* 3L peak and non-peak sequences (gray solid and dotted lines, respectively).

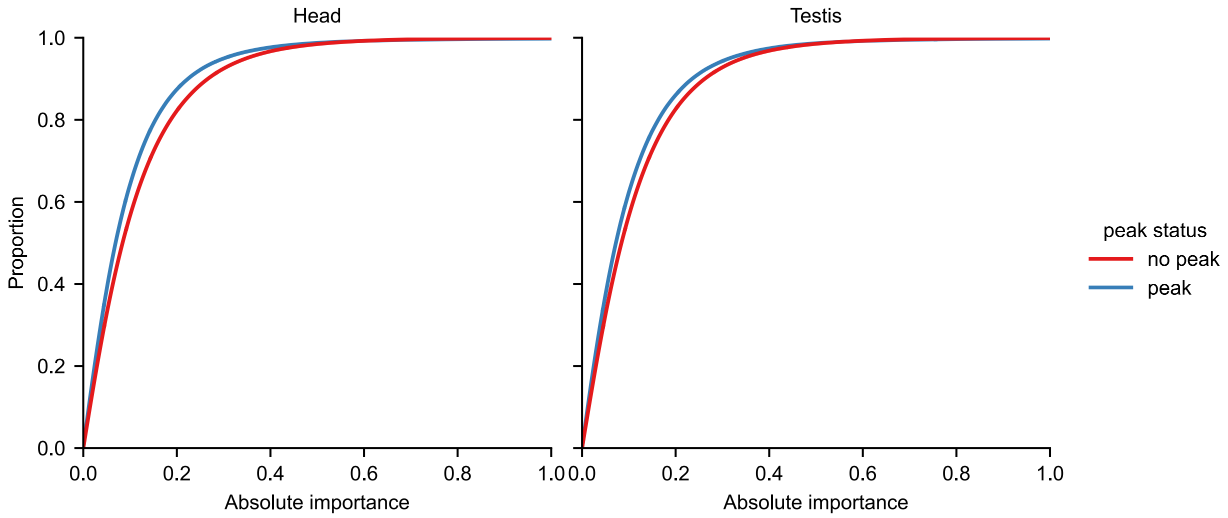


**Supplementary Figure 11. Effect sizes of individual mutations in in silico mutagenesis.** The effect sizes are the absolute importance scores of each site. Only the central 200 bp of each test example was considered.

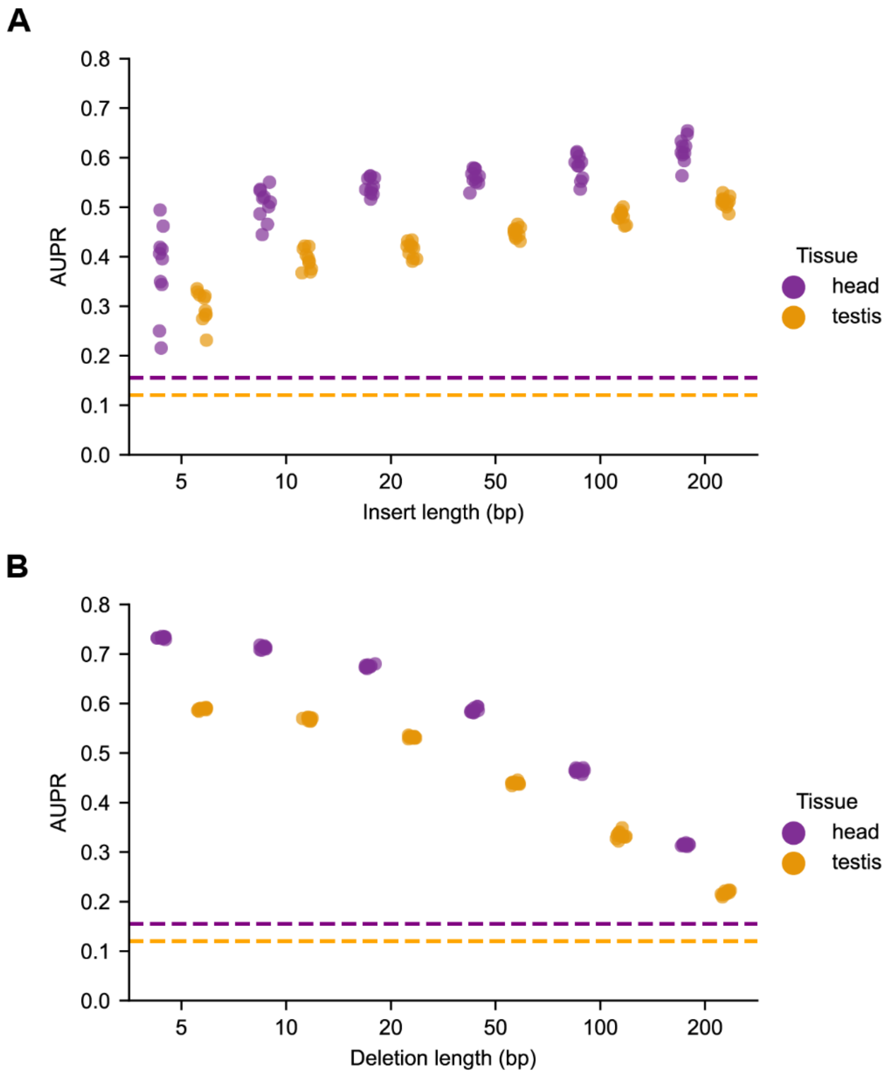


**Supplementary Figure 12. AUPRs for sliding window mutagenesis.** (A) For each example the short subsequence corresponding to the window with largest moving-average importance was inserted into a constant genomic background. Ten sequences with low baseline model outputs were randomly selected, and AUPRs were calculated for model performance. (B) Deletion of windows with highest importance in each example did not substantially diminish model performance for small windows. Deletions were performed by substituting the native sub-sequence with a random sequence, and AUPRs were calculated for model performance. The dotted lines represent baseline (non-predictive) values for AUPR.

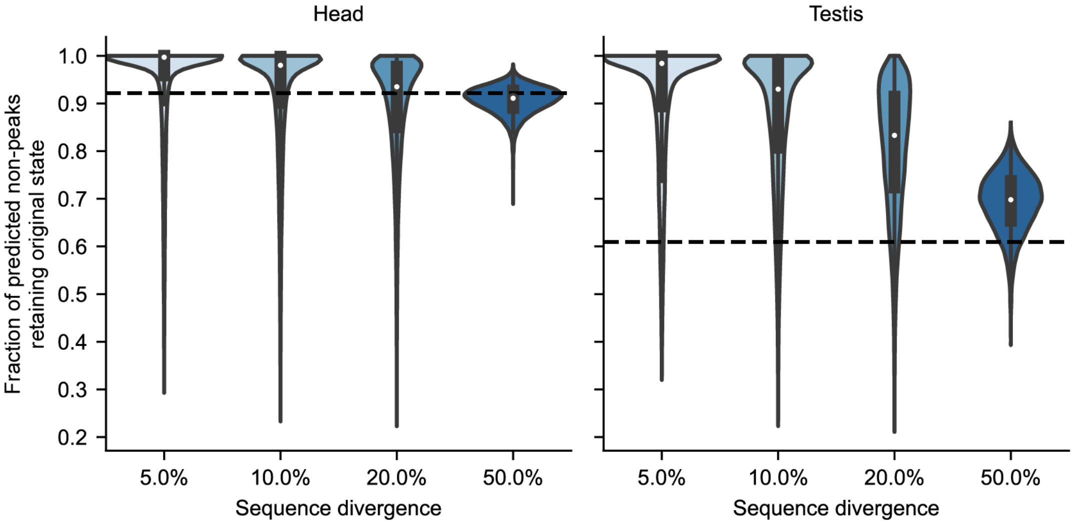


**Supplementary Figure 13. Predicted chromatin accessibility changes as a result of deep mutation.** At each mutational level, 1000 experiments were run for each example, and the fraction of experiments in which a non-peak was predicted to remain a non-peak was calculated. The violin plots show the distributions of these fractions across all non-peak examples. The dotted line represents the fraction of experiments in which a completely random sequence was classified as a non-peak.

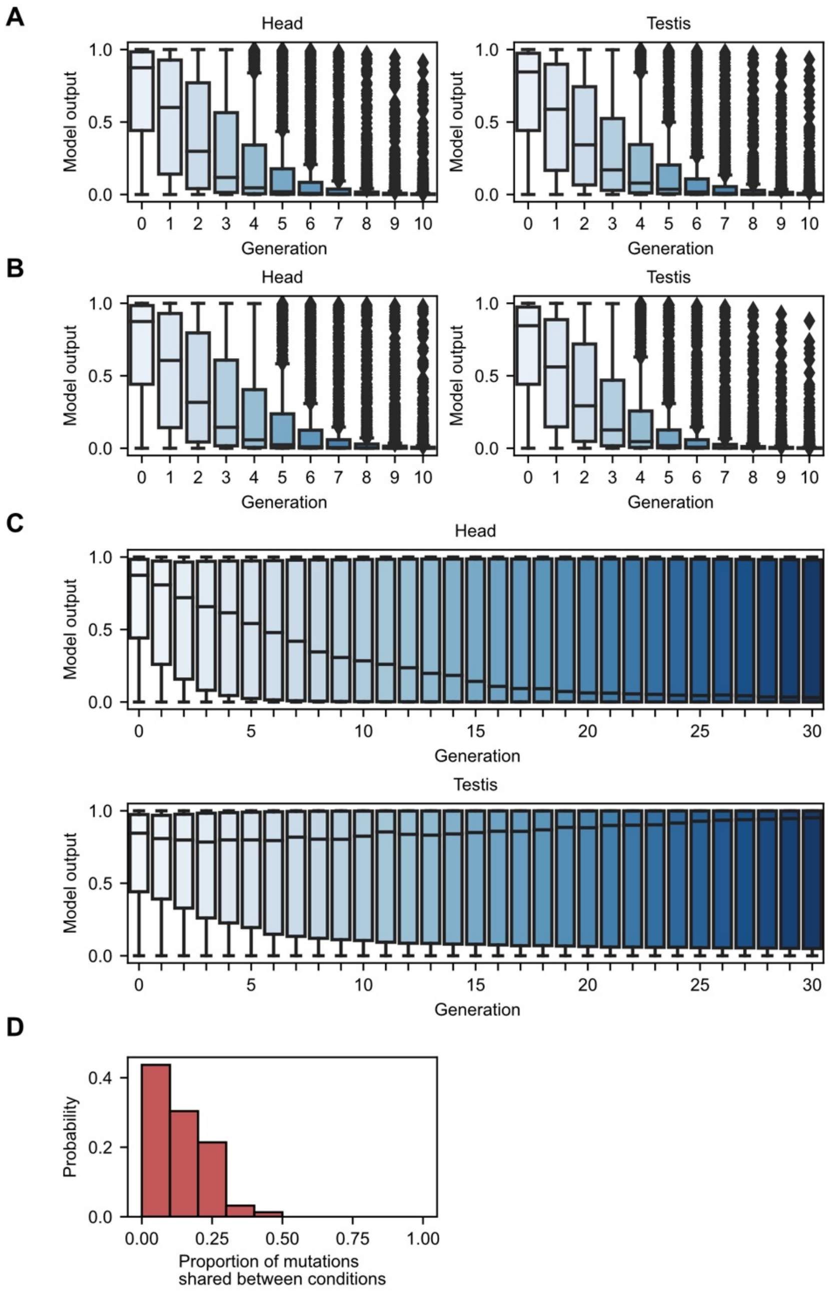


**Supplementary Figure 14. Strong-selection weak-mutation experiments on accessible sequences.** A) Strong selection for inaccessibility in head tissue on 1000 randomly selected peak sequences (peaks in both tissues) results in rapidly decreasing accessibility in both head and testis tissue. B) Using the same examples as in (A), strong selection for inaccessibility in both tissues results in rapidly decreasing accessibility in both tissues. C) Using the same examples as in (A), strong selection for inaccessibility in head and accessibility in testis results in some tissue-specific accessibility, however adaptation is dramatically slowed compared to (A). (D) The proportion of mutations made to examples in (A) also made in (C), revealing distinct mutational paths to accessibility in head tissue.
