## supplementary file 2 for "The evolution and mutational robustness of chromatin accessibility in *Drosophila*": metacluster_list.rtf

motif_category = {'inc_head': ['metacluster_1', 'metacluster_4'], 
                  'dec_head': ['metacluster_0', ], 
                  'inc_testis': ['metacluster_1', 'metacluster_3'],
                  'dec_testis': ['metacluster_0', 'metacluster_2']}


metacluster_0: decrease head, decrease testis
metacluster_1: increase head, increase testis
metacluster_2: no change head, decrease testis
metacluster_3: no change head, increase testis
metacluster_4: increase head, no change testis
