## supplementary file 2 for "The evolution and mutational robustness of chromatin accessibility in *Drosophila*": readme.txt

meme_file_multitask.txt  -- position probability matrices of each motif
metacluster_list.rtf     -- identifies metaclusters from TF-MoDISco
tomtom_jasparinsects.tsv -- results of search from TomTom on Jaspar insects db
